# Can pyrethroid-piperonyl butoxide (PBO) nets reduce the efficacy of indoor residual spraying with pirimiphos-methyl against pyrethroid-resistant malaria vectors?

**DOI:** 10.1101/2021.12.08.471747

**Authors:** Thomas Syme, Martial Gbegbo, Dorothy Obuobi, Augustin Fongnikin, Abel Agbevo, Damien Todjinou, Corine Ngufor

## Abstract

As the uptake of pyrethroid-PBO ITNs increases, their combination with IRS insecticides could become an operational reality in many malaria-endemic communities. Pirimiphos-methyl is a pro-insecticide requiring activation by mosquito cytochrome P450 enzymes to induce toxicity while PBO blocks activation of these enzymes in pyrethroid-resistant vector mosquitoes. PBO may thus antagonise the toxicity of pirimiphos-methyl IRS when combined with pyrethroid-PBO ITNs. The impact of combining two major brands of pyrethroid-PBO ITNs (Olyset^®^ Plus, PermaNet^®^ 3.0) with pirimiphos-methyl IRS (Actellic^®^ 300CS) was evaluated against pyrethroid-resistant *Anopheles gambiae* sl in two parallel experimental hut trials in southern Benin in comparison to bendiocarb IRS and each intervention alone. The wild vector population was resistant to pyrethroids but susceptible to pirimiphos-methyl and bendiocarb. PBO pre-exposure partially restored deltamethrin toxicity but not permethrin. Mosquito mortality in experimental huts was significantly improved in the combinations of bendiocarb IRS with Olyset^®^ Plus (33%) and PermaNet^®^ 3.0 (38%) compared to bendiocarb IRS alone (14–16%, p<0.001), demonstrating an additive effect. Conversely, mortality was significantly reduced in the combinations of pirimiphos-methyl IRS with Olyset^®^ Plus (59%) and PermaNet^®^ 3.0 (55%) compared to pirimiphos-methyl IRS alone (77–78%, p<0.001), demonstrating an antagonistic effect. Combining pirimiphos-methyl IRS with the pyrethroid-PBO ITNs provided significantly improved mosquito mortality (55-59%) compared to the pyrethroid-PBO ITNs alone (22-26%) and improved blood-feeding inhibition relative to the IRS alone. This study provided evidence of an antagonistic effect when pyrethroid-PBO ITNs were combined with pirimiphos-methyl IRS in the same household resulting in lower levels of vector mosquito mortality compared to the IRS alone. Pirimiphos-methyl IRS also showed potential to significantly enhance malaria control when deployed to complement pyrethroid-PBO ITNs in an area where PBO fails to fully restore susceptibility to pyrethroids.

## Introduction

The large-scale implementation of long-lasting insecticidal nets (LLINs), and indoor residual spraying (IRS) has resulted in profound reductions in malaria-associated morbidity and mortality across sub-Saharan Africa, over the last two decades [1]. Unfortunately, resistance to the insecticides applied through these interventions, especially the pyrethroids, is now pervasive in vector populations in malaria-endemic countries [2], threatening to undermine the impact of these interventions. In response, a new generation of novel LLINs and IRS based on new active ingredients with the potential to sustain vector control impact in the face of increasing resistance, have been developed [3]. This includes dual insecticide-treated nets containing a pyrethroid and an alternative effective new compound as well as new IRS insecticides with novel modes of action or improved formulations that have shown potential to provide enhanced control of insecticide-resistant malaria vector populations.

Insecticide-treated nets (ITNs) combining a pyrethroid and piperonyl butoxide (PBO) were the first novel class of ITNs to be developed for malaria vector control. PBO is a synergist that can enhance the impact of pyrethroids and other insecticides by inhibiting metabolic detoxification enzymes associated with resistance, namely cytochrome P450 monooxygenases [4]. These nets received a conditional endorsement from the World Health Organisation (WHO) in 2017 based on results from a cluster-randomised controlled trial in Tanzania demonstrating a reduction in malaria prevalence in communities allocated to pyrethroid-PBO ITNs relative to pyrethroid-only ITNs [5, 6]. A full policy recommendation is now expected after results from a second CRT in Uganda also showed that two brands of pyrethroid-PBO ITNs reduced malaria prevalence relative to pyrethroid-only nets [7]. The WHO endorsement and expanding evidence base for the public health value of pyrethroid-PBO ITNs has prompted mass procurement of these nets by international malaria control agencies [8, 9]. The proportion of pyrethroid-PBO ITNs of all ITNs delivered in sub-Saharan Africa has consequently risen from 3% in 2018 to 35% in 2021 [10]; pyrethroid-PBO ITNs are therefore replacing pyrethroid-only nets in many malaria-endemic countries.

The increasing distribution and intensity of pyrethroid resistance has also affected malaria control policy regarding insecticide choice for IRS over the last decade. To improve vector control impact and preserve pyrethroids for ITNs, African IRS programmes partly suspended the use of pyrethroids and organochlorines in favour of carbamates and organophosphates [11, 12]. Whilst both insecticides showed high toxicity against malaria vectors, the short residual duration of the initial formulations approved for IRS proved prohibitive, necessitating the development of longer-lasting formulations [13]. A new microencapsulated formulation of pirimiphos-methyl was later developed (Actellic^®^ 300CS) demonstrating prolonged activity against pyrethroid-resistant malaria vector mosquitoes, lasting up to 9 months [14, 15]. This formulation subsequently served as the insecticide of choice for the majority of IRS programmes in sub-Saharan Africa [13] providing substantial control of mosquito vectors and malaria across distinct eco-epidemiological settings [6, 16-22].

Where resources are available, there is opportunity to deploy LLINs together with IRS in the same geographical location as a combined intervention approach. The combined LLIN and IRS intervention approach presents opportunities to improve vector control impact by providing multiple chances to target the vector and manage insecticide resistance by presenting two or more insecticides to the vector at the same time. The impact of this approach however depends on several factors including the local vector characteristics and the mode of action of the insecticides involved and interactions that may exist between them [23]. As the uptake of pyrethroid-PBO ITNs increases, their combination with IRS insecticides such as pirimiphos-methyl could become an operational reality in many malaria-endemic communities. Pirimiphos-methyl is a pro-insecticides requiring activation by mosquito enzymes (cytochrome P450 enzymes) to induce toxicity [24]. In mosquitoes with elevated cytochrome P450 enzymes, this activation process will occur faster. By contrast, PBO is combined with pyrethroids on ITNs to enhance the mortality of pyrethroid-resistant vector mosquitoes by inhibiting the activity of these enzymes. It has therefore been hypothesised that the inhibitory action of PBO against cytochrome P450 enzymes may antagonise the toxicity of pro-insecticides like pirimiphos-methyl when used for IRS in combination with pyrethroid-PBO ITNs [25]. Indeed, previous studies on aquatic fleas showed that coadministration of PBO effectively reduced the acute toxicity of organophosphate insecticides requiring metabolic activation by cytochrome P450 but did not affect the toxicity of insecticides not requiring such metabolic activation [26]. Another study assessing the role of PBO in the metabolism of the organophosphorus insecticide, chlorpyrifos in two aquatic invertebrate species also demonstrated that PBO reduced toxicity to the insecticide via cytochrome P450 mediated reactions [27]. Though evidence of this antagonism between PBO and organophosphate insecticides is yet to be demonstrated in bioassays with mosquito vectors of malaria, there have been reports indicating negative interactions in *Anopheles* and *Culex* mosquitoes in bioassays with PBO and other insecticides used for malaria vector control that require metabolic activation [28-30]. A community randomised controlled trial in Tanzania demonstrated a redundant effect on malaria prevalence when adding pirimiphos-methyl IRS in communities with high coverage of pyrethroid-PBO ITNs however, it is not clear whether there was antagonism between the ITNs and IRS [6]. Given the increasing uptake of pyrethroid-PBO ITNs, controlled empirical studies are required, to demonstrate how they interact with different types of IRS and the potential advantages and disadvantages of combining these interventions against different vector populations. Data generated from such trials can be used to inform malaria control policy in areas where the combined implementation of these interventions is anticipated.

In this study, we evaluated the impact of combining two major types of WHO-prequalified pyrethroid-PBO ITNs that have demonstrated additional public health benefit compared to pyrethroid-only nets (Olyset^®^ Plus and PermaNet^®^ 3.0) with IRS using pirimiphos-methyl (Actellic^®^ 300CS) in comparison to bendiocarb IRS in experimental hut trials in a pyrethroid resistance area of southern Benin. The combinations were also compared to each intervention alone.

## Results

### Study site and experimental hut treatments

Experimental huts are used to assess the capacity of indoor vector control interventions to prevent wild vector mosquito entry and feeding and induce early mosquito exiting and mortality when applied in a human-occupied house, under carefully controlled conditions [31, 32]. The hut trials were conducted at the CREC/LSHTM experimental hut station in Covè, southern Benin (7°14′N2°18′E), situated in a vast area of rice irrigation, which provides extensive and permanent breeding sites for mosquitoes. The rainy season extends from March to October and the dry season from November to February. *Anopheles coluzzii* and *An. gambiae* sensu stricto (s.s.) occur in sympatry, with the latter present at lower densities and predominantly in the dry season. The vector population is susceptible to organophosphates and carbamates but exhibits intense resistance to pyrethroids (200-fold). Molecular genotyping and microarray studies have demonstrated a high frequency of the knockdown resistance L1014F allele (>90%) and overexpression of CYP6P3, an enzyme associated with pyrethroid detoxification [33].

Two experimental hut trials were performed in parallel for 4 months between April and July 2020. Trial 1 assessed the impact of combining Olyset^®^ Plus (Sumitomo Chemical), a permethrin-based pyrethroid-PBO ITN with pirimiphos-methyl IRS while Trial 2 assessed the impact of combining PermaNet^®^ 3.0 (Vestergaard Sarl), a deltamethrin-based pyrethroid-PBO ITN with pirimiphos-methyl IRS.

The following six treatments were tested in each of the experimental hut trials:

**Trial 1:**

1. Untreated net
2. Olyset^®^ Plus (Sumitomo Chemical)
3. Bendiocarb IRS applied at 400 mg/m^2^ (Ficam^®^ 80WP, Bayer)
4. Olyset^®^ Plus + Bendiocarb IRS applied at 400 mg/m^2^
5. Pirimiphos-methyl IRS applied at 1000 mg/m^2^ (Actellic^®^ 300CS, Syngenta)
6. Olyset^®^ Plus + Pirimiphos-methyl IRS applied at 1000 mg/m^2^

**Trial 2:**

1. Untreated net
2. PermaNet^®^ 3.0 (Vestergaard Sarl)
3. Bendiocarb IRS applied at 400 mg/m^2^ (Ficam^®^ 80WP, Bayer)
4. PermaNet^®^ 3.0 + Bendiocarb IRS applied at 400 mg/m^2^
5. Pirimiphos-methyl IRS applied at 1000 mg/m^2^ (Actellic^®^ 300CS, Syngenta)
6. PermaNet^®^ 3.0 + Pirimiphos-methyl IRS applied at 1000 mg/m^2^

Olyset^®^ Plus and PermaNet^®^ 3.0 are WHO prequalified pyrethroid-PBO ITNs [3]. Olyset^®^ Plus is made of polyethylene filaments coated with 20 g/Kg of permethrin and 10 g/Kg of PBO. PermaNet^®^ 3.0 consists of polyester side panels coated with deltamethrin at 2.1 g/kg and a polyethylene roof panel incorporating deltamethrin and PBO at 4.0 g/kg and 25 g/kg respectively.

### Susceptibility of wild vector mosquitoes at Covè to insecticides

WHO susceptibility bioassays were conducted in parallel to the experimental hut trial to determine the susceptibility of the vector population at the Covè hut site to the constituent insecticides of the experimental hut treatments. Mortality rates of F1 progeny of field-collected *An. gambiae* s.l. from the Covè hut station following exposure to discriminating doses of deltamethrin and permethrin in WHO cylinder bioassays were low (42% and 11% respectively), confirming a high frequency of pyrethroid resistance in the Covè vector population (Table 1). Pre-exposure to PBO significantly improved mortality with deltamethrin (42% vs. 72%) but not with permethrin (11% vs. 8%). Mortality rates with the discriminating doses of bendiocarb and pirimiphos-methyl were 96% and 99% respectively demonstrating susceptibility to both insecticides. All insecticides induced 100% mortality with the laboratory-maintained, insecticide-susceptible *An. gambiae* s.s. Kisumu strain. No mortality was recorded in the control with either strain.

**Table 1:** WHO susceptibility bioassay results with *Anopheles gambiae* sensu lato from Covè. *Mosquitoes were exposed to the discriminating doses of deltamethrin (0.05%), permethrin (0.75%), bendiocarb (0.1%) and pirimiphos-methyl (0.25%) in four batches of 20–25 per cylinder. The Covè strain was compared with the laboratory-maintained, susceptible Anopheles gambiae sensu stricto Kisumu strain*.

| Treatment | Strain | N exposed | N dead | % Mortality | 95% CIs |
| --- | --- | --- | --- | --- | --- |
| Control | Kisumu | 96 | 0 | 0 | – |
|  | Covè | 98 | 3 | 3 | 0–7 |
| PBO (4%) | Covè | 94 | 0 | 0 | – |
| Deltamethrin (0.05%) | Kisumu | 94 | 94 | 100 | – |
|  | Covè | 90 | 38 | 42 | 32–52 |
| PBO (4%) + Deltamethrin (0.05%) | Covè | 96 | 69 | 72 | 63–81 |
| Permethrin (0.75%) | Kisumu | 91 | 91 | 100 | – |
|  | Covè | 85 | 9 | 11 | 4–17 |
| PBO (4%) + Permethrin (0.75%) | Covè | 85 | 7 | 8 | 2–14 |
| Bendiocarb (0.1%) | Kisumu | 88 | 88 | 100 | – |
|  | Covè | 94 | 90 | 96 | 92–100 |
| Pirimiphos-methyl (0.25%) | Kisumu | 98 | 98 | 100 | – |
|  | Covè | 100 | 99 | 99 | 97–100 |

## Experimental hut results

### Combining pirimiphos-methyl IRS with pyrethroid-PBO ITNs reduces mosquito entry and enhances early exiting

A total of 5,404 wild female *An. gambiae* s.l. were collected in the experimental huts over the 4-month trials (Table 2). In both trials, mosquito entry in huts with the pyrethroid-PBO ITNs alone and IRS alone did not differ significantly from the controls (p>0.05) but was significantly reduced with the pyrethroid-PBO ITN plus IRS combinations compared to the single treatments (p<0.01). Mosquito entry rates did not also differ between the combinations of the pyrethroid-PBO ITN with bendiocarb IRS relative to the combinations with pirimiphos-methyl IRS (p<0.05). Nevertheless, IRS treatments could not be rotated and thus, treatment-induced deterrence cannot be fully distinguished from differential attractiveness due to hut position.

**Table 2:** Entry and exiting of wild, pyrethroid-resistant *Anopheles gambiae* sensu lato in experimental huts in Covè, southern Benin treated with pyrethroid-PBO ITNs and pirimiphos-methyl IRS applied alone and in combination. *Results are presented separately for the trials involving Olyset^®^ Plus (Trial 1) and PermaNet^®^ 3.0 (Trial 2)*.

| Treatment | Total females caught* | % Deterrence | Total exiting | % Exophily* | 95% CIs |
| --- | --- | --- | --- | --- | --- |
| **Trial 1** |  |  |  |  |  |
| Untreated net | 664<sup>a</sup> | – | 241 | 36<sup>a</sup> | 33–40 |
| Olyset Plus | 511<sup>a</sup> | 23 | 390 | 76<sup>b</sup> | 73–80 |
| Bendiocarb IRS | 556<sup>a</sup> | 16 | 282 | 51<sup>c</sup> | 47–55 |
| Olyset Plus + Bendiocarb IRS | 288<sup>b</sup> | 57 | 228 | 79<sup>b</sup> | 75–84 |
| P-methyl IRS | 531<sup>a</sup> | 20 | 281 | 53<sup>c</sup> | 49–57 |
| Olyset Plus + P-methyl IRS | 304<sup>b</sup> | 54 | 270 | 89<sup>d</sup> | 85–92 |
| **Trial 2** |  |  |  |  |  |
| Untreated net | 581<sup>u</sup> | – | 226 | 39<sup>u</sup> | 35–43 |
| PermaNet 3.0 | 488<sup>u</sup> | 16 | 369 | 76<sup>v</sup> | 72–79 |
| Bendiocarb IRS | 575<sup>u</sup> | 1 | 360 | 63<sup>w</sup> | 59–67 |
| PermaNet 3.0 + Bendiocarb IRS | 233<sup>v</sup> | 60 | 185 | 79<sup>v</sup> | 74–85 |
| P-methyl IRS | 450<sup>u</sup> | 23 | 311 | 69<sup>x</sup> | 65–73 |
| PermaNet 3.0 + P-methyl IRS | 223<sup>v</sup> | 62 | 195 | 87<sup>y</sup> | 83–92 |

\*For each trial, values on this column sharing a superscript letter do not differ significantly.

The proportion of mosquitoes exiting into the veranda of the huts with the untreated net controls was 36% and 39% for Trials 1 and 2 respectively. Exiting rates were higher with the pyrethroid-PBO ITNs alone (76%) relative to the IRS insecticides alone (51-63% with bendiocarb and 53-69% with pirimiphos-methyl IRS, P<0.005). In both trials, the highest levels of mosquito exiting were achieved with the pyrethroid-PBO ITN plus IRS combinations (79% with bendiocarb and 87-89% with pirimiphos-methyl IRS). Between the combinations, adding pirimiphos-methyl IRS to the pyrethroid-PBO ITN provided significantly higher levels of mosquito exiting relative to adding bendiocarb IRS (87-89% vs. 79%, P<0.05).

### Combining pirimiphos-methyl IRS with pyrethroid-PBO ITNs reduces mosquito feeding relative to the IRS alone

Blood-feeding rates in huts with the untreated net controls were 72% and 67% for Trials 1 and 2 respectively (Table 3). The IRS treatments did not provide any blood-feeding inhibition relative to the controls. Blood-feeding inhibition rates were high with the pyrethroid-PBO ITN plus IRS combinations in both trials (69-79% with Olyset^®^ Plus and 63-66% with PermaNet^®^ 3.0) and this was generally similar to what was observed with the pyrethroid-PBO ITNs alone (67% with Olyset^®^ Plus and 65% with PermaNet^®^) showing that the high levels of blood-feeding inhibition in the combinations, was mostly due to the ITNs. Blood-feeding inhibition with the PermaNet^®^ 3.0 plus pirimiphos-methyl IRS combination (66%) was similar to the PermaNet^®^ 3.0 plus bendiocarb IRS combination (63%, p=0.71) meanwhile the Olyset^®^ Plus and pirimiphos-methyl IRS combination induced higher levels of blood-feeding inhibition compared to the Olyset^®^ Plus and bendiocarb IRS combination (79% vs. 69%, p=0.036). Personal protection levels showed a similar trend and were also higher with the combinations (86-90%) relative to the IRS alone (−33 −13%)

**Table 3:** Blood-feeding results of wild, pyrethroid-resistant *Anopheles gambiae* sensu lato exposed to pyrethroid-PBO ITNs and pirimiphos-methyl IRS applied alone and in combination in experimental huts in Covè, southern Benin. *Results are presented separately for the trials involving Olyset^®^ Plus (Trial 1) and PermaNet^®^ 3.0 (Trial 2)*.

| Treatment | Total Females Caught | Total blood-fed | % Blood-feeding* | 95% CIs | % Blood-feeding inhibition | % Personal protection |
| --- | --- | --- | --- | --- | --- | --- |
| **Trial 1** |  |  |  |  |  |  |
| Untreated net | 664 | 481 | 72<sup>a</sup> | 69–76 | – | – |
| Olyset Plus | 511 | 122 | 24<sup>b</sup> | 20–28 | 67 | 75 |
| Bendiocarb IRS | 556 | 528 | 95<sup>c</sup> | 93–97 | -31 | -10 |
| Olyset Plus + Bendiocarb IRS | 288 | 64 | 22<sup>b</sup> | 17–27 | 69 | 87 |
| P-methyl IRS | 531 | 420 | 79<sup>d</sup> | 76–83 | -9 | 13 |
| Olyset Plus + P-methyl IRS | 304 | 47 | 16<sup>e</sup> | 11–20 | 79 | 90 |
| **Trial 2** |  |  |  |  |  |  |
| Untreated net | 581 | 391 | 67<sup>u</sup> | 64–71 | – | – |
| PermaNet 3.0 | 488 | 115 | 24<sup>v</sup> | 20–27 | 65 | 71 |
| Bendiocarb IRS | 575 | 519 | 90<sup>w</sup> | 88–93 | -34 | -33 |
| PermaNet 3.0 + Bendiocarb IRS | 233 | 54 | 23<sup>v</sup> | 18–29 | 66 | 86 |
| P-methyl IRS | 450 | 406 | 90<sup>w</sup> | 88–93 | -34 | -4 |
| PermaNet 3.0 + P-methyl IRS | 223 | 55 | 25<sup>v</sup> | 19–30 | 63 | 86 |

\*For each trial, values on this column sharing a superscript letter do not differ significantly.

### Combining pirimiphos-methyl IRS with pyrethroid-PBO ITNs reduces mosquito mortality relative to the IRS alone

Wild vector mosquito mortality in the controls was 3% in Trial 1 and 2% in Trial 2 (Table 4). The pyrethroid-PBO ITNs killed relatively low mosquito proportions (22% with Olyset^®^ Plus and 26% with PermaNet^®^ 3.0) though this was higher than what was observed with bendiocarb IRS alone (14 in Trial 1 and 16% in Trial 2, P<0.01). The highest mortality was achieved with pirimiphos-methyl IRS alone (77% in Trial 1 and 78% in Trial 2). In both trials, mortality in the pyrethroid-PBO ITN plus pirimiphos-methyl IRS combinations was significantly reduced compared to pirimiphos-methyl IRS alone (77% with pirimiphos-methyl IRS vs. 59% with Olyset^®^ Plus plus pirimiphos-methyl IRS, p<0.001 and 78% with pirimiphos-methyl IRS vs. 55% with PermaNet^®^ 3.0 plus pirimiphos-methyl IRS, p<0.001), demonstrating an antagonistic effect. Conversely, mortality was significantly higher in the combinations of bendiocarb IRS with Olyset^®^ Plus (33%) and PermaNet^®^ 3.0 (38%) than bendiocarb IRS alone (14–16%, p<0.001), demonstrating an additive effect (Table 4). Nevertheless, both pyrethroid-PBO plus IRS combinations induced significantly higher mortality compared to the pyrethroid-PBO ITNs alone (p<0.001). Between the combinations, mortality was consistently higher with the pyrethroid-PBO ITN plus pirimiphos-methyl IRS combinations (55-59%) compared to the pyrethroid-PBO ITN plus bendiocarb IRS (33-39%, p<0.001).

**Table 4:** Mortality results of wild, pyrethroid-resistant *Anopheles gambiae* sensu lato in experimental huts in Covè, southern Benin treated with pyrethroid-PBO ITNs, and pirimiphos-methyl IRS applied alone and in combination. *Results are presented separately for the trials involving Olyset^®^ Plus (Trial 1) and PermaNet^®^ 3.0 (Trial 2)*.

| Treatment | Total Females Caught | Total dead | % Mortality* | 95% CIs |
| --- | --- | --- | --- | --- |
| **Trial 1** |  |  |  |  |
| Untreated net | 664 | 22 | 3<sup>a</sup> | 2–5 |
| Olyset Plus | 511 | 114 | 22<sup>b</sup> | 19–26 |
| Bendiocarb IRS | 556 | 87 | 16<sup>c</sup> | 13–19 |
| Olyset Plus + Bendiocarb IRS | 288 | 95 | 33<sup>d</sup> | 28–38 |
| P-methyl IRS | 531 | 411 | 77<sup>e</sup> | 74–81 |
| Olyset Plus + P-methyl IRS | 304 | 178 | 59<sup>f</sup> | 53–64 |
| **Trial 2** |  |  |  |  |
| Untreated net | 581 | 12 | 2<sup>u</sup> | 1–3 |
| PermaNet 3.0 | 488 | 127 | 26<sup>v</sup> | 22–30 |
| Bendiocarb IRS | 575 | 80 | 14<sup>w</sup> | 11–17 |
| PermaNet 3.0 + Bendiocarb IRS | 233 | 89 | 38<sup>x</sup> | 32–44 |
| P-methyl IRS | 450 | 350 | 78<sup>y</sup> | 74–82 |
| PermaNet 3.0 + P-methyl IRS | 223 | 122 | 55<sup>z</sup> | 48–61 |

\*For each trial, values on this column sharing a superscript letter do not differ significantly.

The monthly mortality rates of wild vector mosquitoes which entered the experimental huts during the trials are presented in Fig 1 for the combinations with bendiocarb IRS and Fig 2 for the combinations with pirimiphos-methyl IRS. In both trials, mosquito mortality rates in huts with the pyrethroid-PBO ITNs plus bendiocarb IRS declined sharply over time from 65-75% in month 1 to 33-38% in month 3, nevertheless, it was consistently higher than the single treatments alone (Fig 1). Mosquito mortality in the combination of Olyset^®^ Plus plus pirimiphos-methyl IRS was similar to the IRS alone in month 1 (>90%) but declined substantially relative to the IRS in subsequent months. With the PermaNet^®^ 3.0 plus pirimiphos-methyl IRS combination, mosquito mortality was consistently lower than the IRS alone throughout the trial. Through the four months of the trials, the pyrethroid-PBO ITN plus pirimiphos-methyl IRS combinations consistently induced higher levels of mosquito mortality relative to the pyrethroid-PBO ITNs alone.

**Fig 1:**
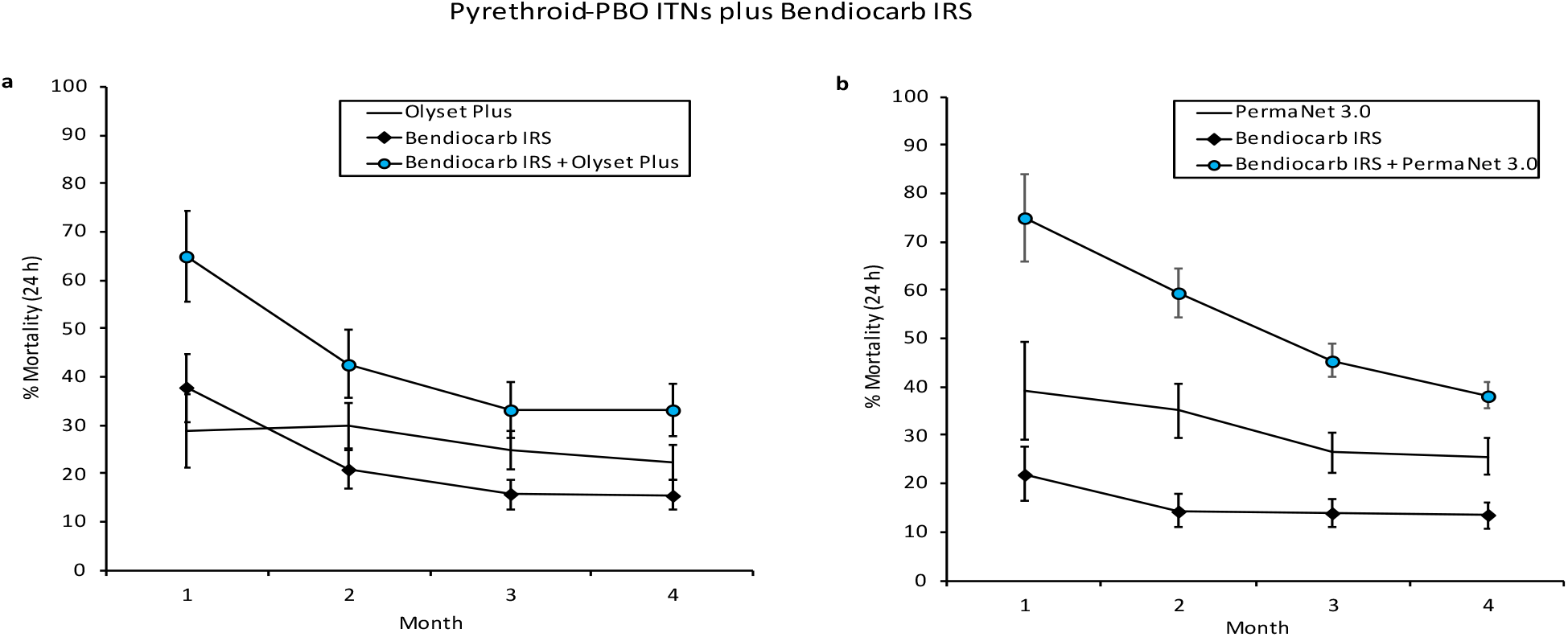
Monthly mortality rates of wild, pyrethroid-resistant *Anopheles gambiae* sensu lato entering experimental huts with pyrethroid-PBO ITNs and bendiocarb IRS, applied alone and in combination in Covè, southern Benin. *Panel a presents results from the trial with Olyset^®^ Plus (Trial 1) and panel b presents results from the trial with PermaNet^®^ 3.0 (Trial 2). Error bars represent 95% CIs. Monthly mortality rates are cumulated with increasing time elapsed from onset of the trial*.

**Fig 2:**
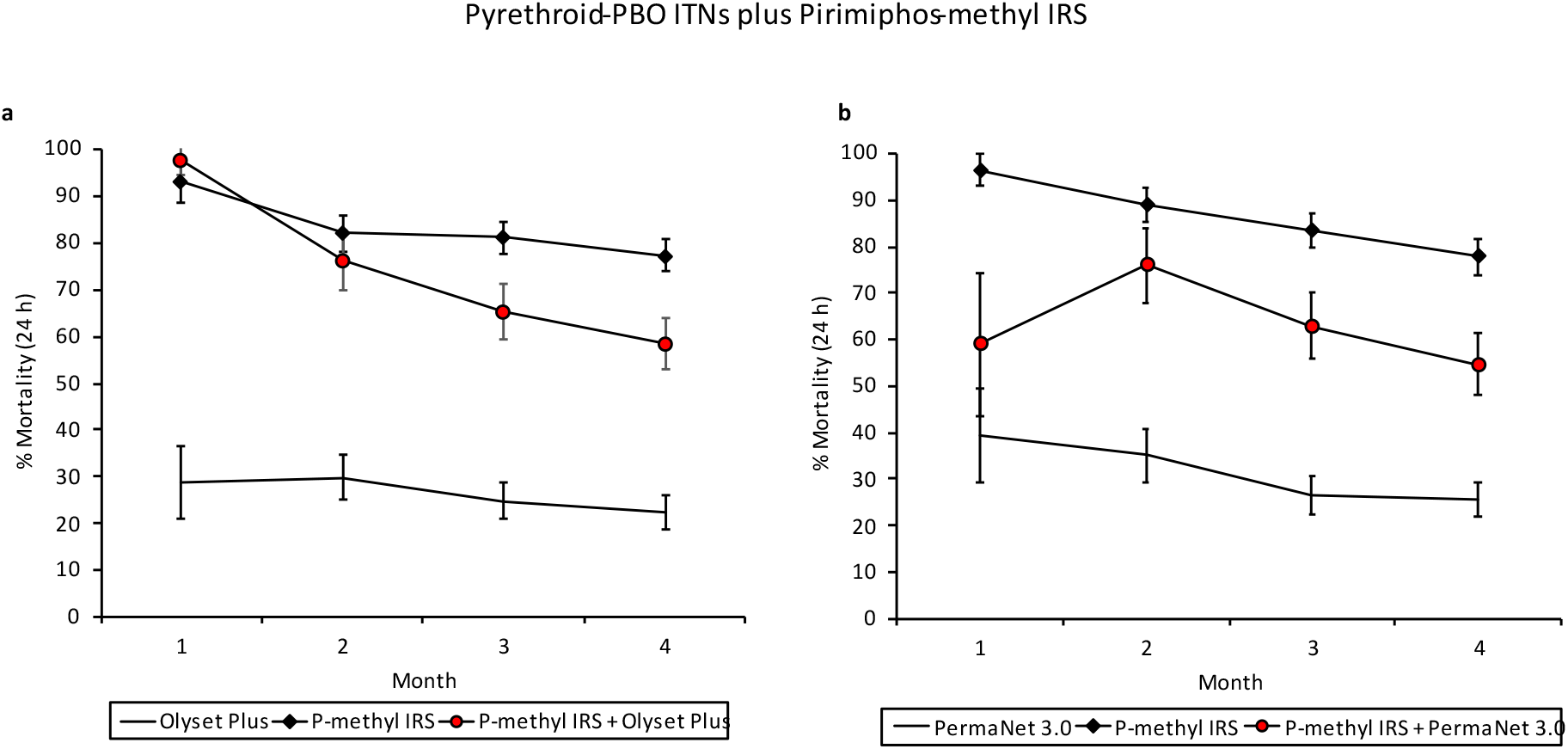
Monthly mortality rates of wild, pyrethroid-resistant *Anopheles gambiae* sensu lato entering experimental huts with pyrethroid-PBO ITNs and pirimiphos-methyl IRS, applied alone and in combination in Covè, southern Benin. *Panel a presents results from the trial with Olyset^®^ Plus and panel b presents results from the trial with PermaNet^®^ 3.0. Error bars represent 95% CIs. Monthly mortality rates are cumulated with increasing time elapsed from the onset of the trial*.

### Tunnel test bioassays

To further assess the efficacy of the pyrethroid-PBO ITNs and help explain the findings in the experimental huts, tunnel tests were performed using the susceptible *An.gambiae* Kisumu strain and pyrethroid-resistant *An. gambiae* sl from Covè on net samples (30×30cm) obtained from Olyset^®^ Plus and PermaNet^®^ 3.0 nets. The tunnel test is an overnight animal baited bioassay that simulates host-seeking behaviour of vector mosquitoes under controlled laboratory conditions. Both pyrethroid-PBO ITNs were compared to pyrethroid-only nets which contained similar pyrethroid-insecticides (Olyset^®^ Net and PermaNet^®^ 2.0). Mortality in the control tunnels was 11% with the susceptible Kisumu strain and 2% with the pyrethroid-resistant Covè strain. All ITN types induced >98% mortality with the susceptible Kisumu strain. Olyset^®^ Plus killed significantly higher proportions of the Covè mosquitoes compared to Olyset^®^ Net (90% vs. 42%). With PermaNet^®^ 3.0, mortality of Covè mosquitoes exposed to netting pieces obtained from the PBO treated roof of the net (68%) was higher compared to net pieces obtained from the sides of the net (34%) and PermaNet^®^ 2.0 (27%). The results, therefore, showed higher levels of mortality against pyrethroid-resistant mosquitoes from the Covè experimental hut station with the pyrethroid-PBO ITNs relative to pyrethroid-only nets. More detailed results from the tunnel tests are available in the supplementary information (Table S1).

### Wall cone bioassays

WHO wall cone bioassays were performed on the IRS treated experimental hut walls 1 week, 2 months and 4 months after application of IRS treatments to assess their residual efficacy with increasing time elapsed from spraying. Mortality rates of the insecticide-susceptible *An. gambiae* s.s. Kisumu strain and pyrethroid-resistant *An. gambiae* s.l. Covè strain following exposure to IRS-treated surfaces in 30 min wall cone bioassays are presented in Fig 4. Mortality of mosquitoes exposed to pirimiphos-methyl treated walls was high (>90%) at all time points with both strains, showing no evidence of a decline in residual activity. In contrast, wall cone bioassay mortality on bendiocarb-treated surfaces declined rapidly, beginning at 67% and 39% in week 1 and falling to lows of 2% and 3% in month 4 with the Kisumu and Covè strains respectively. No mortality was recorded in the controls at any time point with either strain.

**Fig 3:**
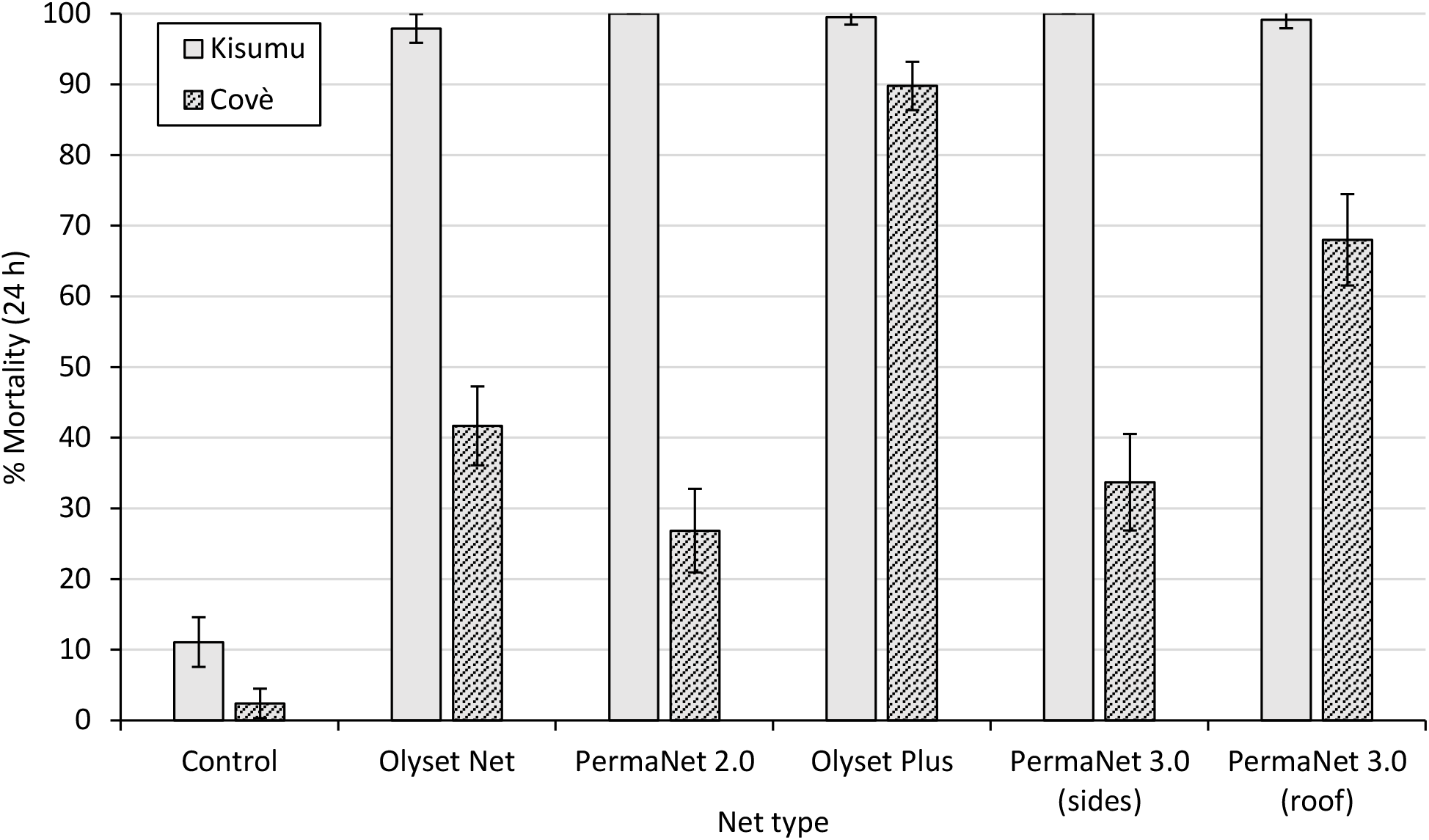
Mortality (24h) of susceptible *Anopheles gambiae* Kisumu and pyrethroid-resistant An gambiae sl Covè strains exposed to Olyset^®^ Plus and PermaNet^®^ 3.0 in tunnel tests. *Error bars represent 95% CIs. Both pyrethroid-PBO nets were compared to pyrethroid-only nets (Olyset^®^ Net and PermaNet^®^ 2.0). PermaNet^®^ 3.0 contains PBO only on the roof of the net. With both strains, 80– 120 mosquitoes were exposed overnight to netting pieces cut from whole nets in 2–3 replicate tunnel tests*.

**Fig 4:**
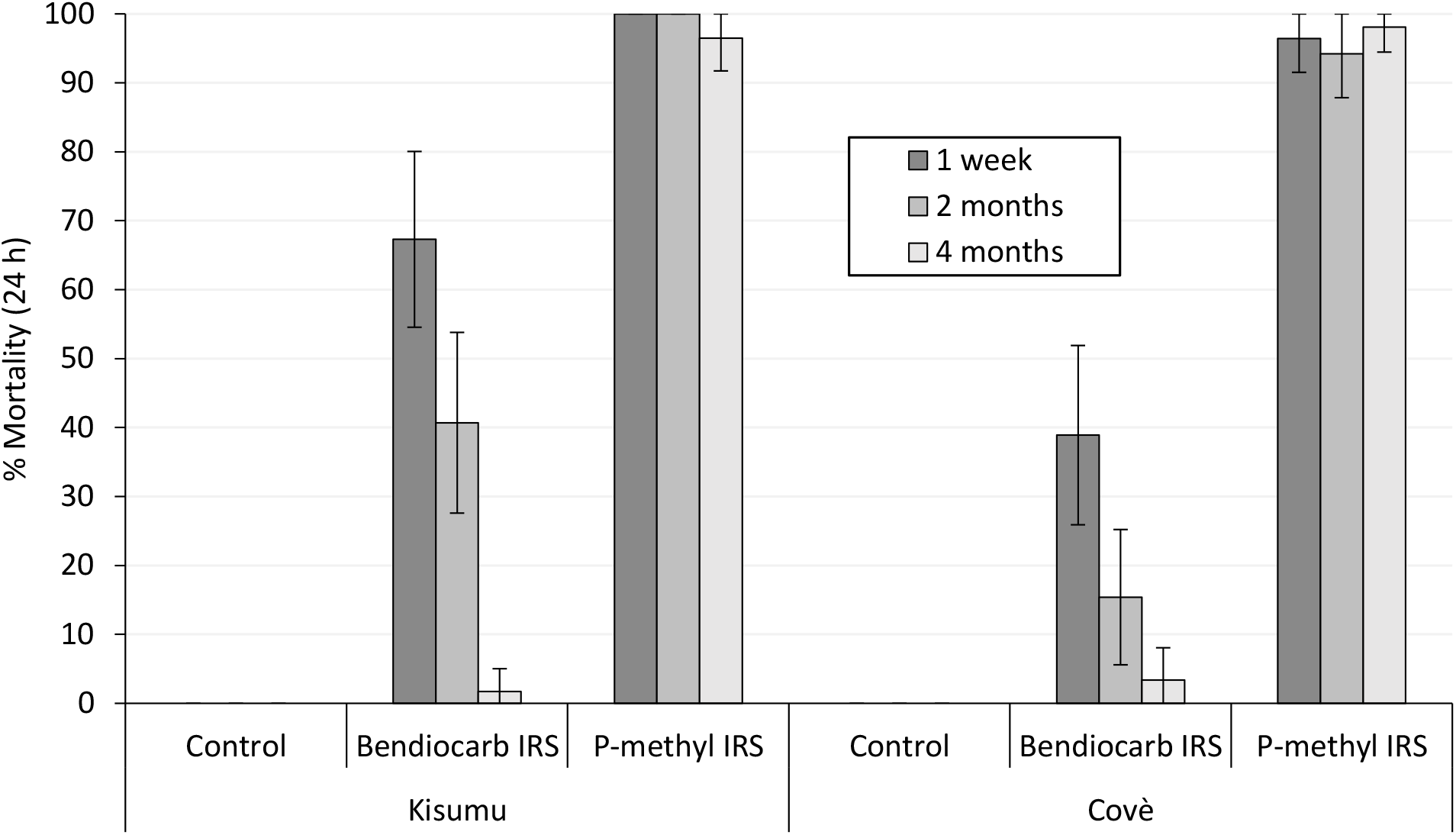
Mortality (24 h) of insecticide-susceptible *Anopheles gambiae* sensu stricto Kisumu strain and pyrethroid-resistant *Anopheles gambiae* sensu lato Covè strain exposed to treated experimental hut walls in 30 min wall cone bioassays. Error bars represent 95% CIs. Approximately 50, 3–5-day old mosquitoes of each strain were exposed to treated walls for 30 mins in 5 batches of 10, 1 week 2 months and 4 months after application of treatments. Mortality was recorded after 24 h.

## Discussion

As the uptake of pyrethroid-PBO ITNs continues to rise, their combined use with IRS could become an operational reality in many malaria-endemic settings across Africa. Given the inhibitory effect of PBO on mosquito cytochrome P450 enzymes, the WHO had temporarily recommended against the use of pyrethroid-PBO ITN in areas programmed for IRS with pirimiphos-methyl IRS until further evidence on the potential antagonism between PBO and the organo-thiophosphate pro-insecticide becomes available [25]. In this study, we found evidence of an antagonistic effect when pyrethroid PBO ITNs were combined with pirimiphos-methyl IRS in the same household in a pyrethroid-resistant area in Southern Benin where resistance was partly conferred by over-expression of mosquito cytochrome P450 enzymes [33]. Unlike the combination of the pyrethroid-PBO nets with bendiocarb IRS which provided improved vector mosquito mortality compared to bendiocarb IRS alone, combining these nets with pirimiphos-methyl IRS induced significantly lower levels of vector mosquito mortality rates compared to pirimiphos-methyl IRS alone. This effect was remarkably similar between the two brands of pyrethroid-PBO ITNs tested across both hut trials indicating that the negative interaction of the combination on mosquito mortality was less affected by the design and specifications of the pyrethroid-PBO ITN. Wild vector mosquitoes which entered the huts with the combined treatment may have contacted the pirimiphos-methyl IRS on the hut wall only after picking up PBO from the ITN while attempting to blood-feed on the sleeper under the net. This pre-exposure to PBO on the ITN could have prevented the metabolic activation of the IRS insecticide in these mosquitoes, resulting in lower mortality rates relative to the IRS alone. This finding supports WHO’s recommendation against the deployment of pyrethroid-PBO ITNs in areas that have already been programmed for IRS with pirimiphos-methyl IRS. This is mostly important from a programmatic perspective where a vector control programme is faced with multiple choice of ITN types to deploy as an additional intervention to enhance vector control impact in an area that is already dedicated to IRS with pirimiphos-methyl. In such a scenario, other types of ITNs like pyrethroid-only nets which were recently shown to complement pirimiphos-methyl IRS when applied together [23], should be considered.

These findings should however not be interpreted to mean that pirimiphos-methyl IRS must not be deployed to complement pyrethroid-PBO ITNs. Though pyrethroid-PBO ITNs have consistently shown improved performance in experimental hut trials against pyrethroid-resistant malaria vectors compared to pyrethroid-only nets [34], the margin appears to vary depending on the intensity and mechanisms of pyrethroid-resistance encountered. In our study, the pyrethroid-PBO ITNs when applied alone in a hut induced low vector mosquito mortality (22-26%) compared to what has been observed in hut trials against the same vector population with another type of novel dual ITN (71-76%) [35, 36]. Similar levels of hut mortality rates with pyrethroid-PBO ITNs (<30%) and failure of PBO to fully restore pyrethroid susceptibility in bioassays has also been reported from studies in Burkina Faso [37], Cameroon [38], Côte d’lvoire [39] and Senegal [40]. This may indicate the presence of complex resistance mechanisms in the West African region unaffected by PBO. Compared to East Africa, West Africa has shown historically higher intensity of pyrethroid resistance in malaria vectors which continues to increase over time [41], yet the public health value of pyrethroid-PBO ITNs has not been assessed in the region. It is therefore unclear whether these nets will provide the same improved epidemiological impact over pyrethroid-only nets in West Africa as observed in the East African community trials which have been the basis of their endorsement for malaria control. Our results showed a significant improvement in mosquito mortality when pirimiphos-methyl IRS was combined with the pyrethroid-PBO ITNs (55-59%) compared to the pyrethroid-PBO ITNs alone (22-26%). Mosquito entry and feeding rates were also significantly lower with the combination compared to the pyrethroid-PBO ITNs alone. This provides some justification for deploying pirimiphos-methyl IRS to complement pyrethroid-PBO nets in an area of high and complex pyrethroid-resistance where local vectors are less susceptible to the synergistic effect of PBO. Due to the limited choice of insecticides available for IRS, where a beneficial effect of the combination compared to the pyrethroid-PBO ITN alone has been established, pirimiphos-methyl IRS should continue to be used even in the presence of high coverage with pyrethroid-PBO ITNs, preferably as part of an IRS rotational strategy. In line with WHO recommendations for insecticide resistance management [42], the sustained rotation of multiple insecticide modes of action for IRS may help preserve efficacy to insecticides approved for IRS.

Considering the significant increase in mosquito mortality with the combination compared to the pyrethroid-PBO ITN alone in our study, it is unclear whether the redundant effect of adding pirimiphos-methyl IRS to pyrethroid-PBO ITNs on clinical malaria observed in the community randomised trial in Tanzania [6] applies to other settings. Unlike our study which showed no restoration of susceptibility to permethrin in bioassays following pre-exposure to PBO, restoration of susceptibility to permethrin - the pyrethroid insecticide used on the pyrethroid-PBO ITN tested in the Tanzanian trial (Olyset^®^ Plus) - was substantially high in the vector population from the study area (from 18% to 94%) [43]. Hence, the level of control achieved with pyrethroid-PBO ITN alone in the Tanzanian trial may have been optimal making the addition of pirimiphos-methyl IRS unnecessary. This may not be the case in many communities in West Africa considering the afore-mentioned low levels of pyrethroid-PBO synergism reported in susceptibility bioassays and hut trials conducted across the region; in contrast, the addition of pirimiphos-methyl IRS to pyrethroid-PBO ITNs in these communities could be more beneficial for control of clinical malaria compared to the ITN alone. As the range of vector control products available to vector control programmes expands, the choice of interventions and product brands must be aligned to local contexts and guided by local evidence. To help guide vector control policy, epidemiological trials and/or other empirical studies investigating the impact and cost-effectiveness of pyrethroid-PBO nets in communities in the West African region as well as their combination with IRS using pro-insecticides like pirimiphos-methyl will be necessary.

Although the inhibitory effect of PBO on mosquito cytochrome P450 enzymes formed the basis of our hypothesis for the antagonism observed between pirimiphos-methyl IRS and pyrethroid-PBO ITNs in the experimental huts, behavioural interactions could also have contributed. In both trials, mosquito exiting rates into the veranda traps and blood-feeding inhibition were consistently higher in the combinations compared to the IRS insecticides alone. This could be attributed to the excito-repellent property of the pyrethroid in the ITN stimulating directed movement of mosquitoes away from their source [44]. This early exiting effect was however significantly higher in the combination with pirimiphos-methyl IRS compared to the combination with bendiocarb IRS suggesting a behavioural interaction in the presence of pirimiphos-methyl that may have driven more mosquitoes to exit into the veranda, thus reducing mosquitoe’s contact with pirimiphos-methyl IRS treated walls. This may have compromised the impact of the pirimiphos-methyl IRS in the combination compared to when applied alone and to the combination with bendiocarb. Further studies to assess mosquito flight behaviour, in the presence of the combinations would provide useful insight into behavioural interactions between the treatments. Controlled laboratory assays which minimise behavioural responses and assess P450 enzyme activity in exposed mosquitoes, could also elucidate the potential role of P450 enzyme inhibition in the antagonism observed in the experimental huts.

While our trial focused on investigating the impact of combining pyrethroid-PBO ITNs with pirimiphos-methyl IRS in the same households, there is a paucity of information on the interactions between these nets and other newly developed vector control products which contain new public health insecticides such as the neonicotinoid, clothianidin and the pyrrole chlorfenapyr. Chlorfenapyr, an insecticide used on ITNs [35] and being considered for IRS [45, 46], also requires activation by mosquito P450 enzymes [47]. Laboratory experiments have indicated the potential of PBO to antagonise the toxicity of chlorfenapyr against mosquitoes in bioassays [29, 30]. Studies investigating the impact of co-deploying pyrethroid-PBO ITNs together with pyrethroid-chlorfenapyr ITNs or chlorfenapyr IRS in the same household will be essential to help inform optimal co-deployment policy.

## Conclusion

Our study provides the first evidence of an antagonistic effect when pyrethroid-PBO ITNs are combined with pirimiphos-methyl IRS in the same household resulting in lower levels of vector mosquito mortality compared to the IRS alone. In line with WHO recommendations, vector control programmes faced with multiple choice of ITN types to deploy as an additional intervention to improve vector control impact in an area dedicated to IRS with pirimiphos-methyl, may consider other types of ITNs like pyrethroid-ITNs which can better complement pirimiphos-methyl IRS when deployed together. Nevertheless, the pyrethroid-PBO ITNs performed poorly probably due to the lower levels of restoration of pyrethroid susceptibility with PBO in the vector population. Combining these nets with pirimiphos-methyl IRS provided significantly improved vector control compared to the net alone demonstrating the potential for pirimiphos-methyl IRS to enhance malaria control when deployed to complement pyrethroid-PBO ITNs in areas where PBO fails to fully restore susceptibility to pyrethroids.

## Materials and methods

### WHO susceptibility bioassays

Adult F1 progeny of field-collected *An. gambiae* s.l. from the Covè hut station were exposed to filter papers treated with discriminating doses of deltamethrin (0.05%), permethrin (0.75%), bendiocarb (0.1%) and pirimiphos-methyl (0.25%) in WHO cylinders [48]. Deltamethrin and permethrin were also tested with 60 mins pre-exposure to PBO (4%) to assess the involvement of metabolic enzymes in pyrethroid resistance. A comparison was made with the laboratory-maintained, susceptible *An. gambiae* s.s. Kisumu strain. Approximately 100, 3–5-day old mosquitoes of each strain were exposed to each insecticide for 60 mins in four batches of 20–25. Similar numbers of mosquitoes were concurrently exposed to untreated filter papers as a control. At the end of exposure, mosquitoes were transferred to appropriately labelled holding tubes, provided access to 10% (w/v) glucose solution and held at 27±2°C and 75±10% relative humidity. Knockdown was recorded 60 mins after exposure and delayed mortality after 24 h for all treatments. The insecticide-treated filter papers were obtained from Universiti Sains Malaysia.

### Experimental huts and treatment application

The experimental huts used for the trials were of standard West African design, made from concrete bricks with cement-plastered walls and ceiling and a corrugated iron roof. A wooden-framed veranda projects from the rear wall of each hut to capture mosquitoes exiting due to behavioural or insecticidal effects. Each hut is constructed on a water-filled moat to preclude entry of predators and mosquito entry occurs via four window slits measuring 1 cm positioned on two sides of the hut.

Prior to the application of the IRS treatments, the IRS insecticide formulations were mixed with appropriate volumes of water to obtain insecticide solutions at desired concentrations. Interior hut walls and ceilings were sprayed from top to bottom by a trained spray team using a Hudson X-pert^®^ compression sprayer equipped with a flat-fan nozzle. Spray swaths were pre-marked on hut walls and ceilings using chalk to improve the accuracy of spraying. To assess the quality of spray application, the spray tank was weighed before and after spraying to assess the volume of insecticide solution applied. All volumes of insecticide solution applied were within the ±30% acceptable deviation from the target, indicating that the treatments were applied correctly. Bed nets were erected over sleeping areas by tying the four corners of the roof panel to nails positioned at the four uppermost corners inside the hut. To simulate wear and tear from routine household use, each ITN received 6 holes each measuring 16 cm^2^ cut into each side of the net.

### Experimental hut trial procedure

During the hut trials, volunteer sleepers were rotated between experimental huts daily to mitigate the impact of individual attractiveness whilst bed nets were rotated weekly between the huts to reduce the impact of hut position on mosquito entry. Three (3) replicate bed nets were used per treatment and rotated within the treatment every 2 days. IRS treatments cannot be rotated and thus remained fixed throughout the trial. Consenting human volunteer sleepers slept in experimental huts between 21:00 and 06:00 to attract free-flying mosquitoes. Each morning, volunteer sleepers collected all live and dead mosquitoes from the different compartments of the hut (under the net, room, veranda) using a torch and aspirator and placed them in correspondingly labelled plastic cups. Mosquito collections were then transferred to the field laboratory for morphological identification using taxonomic keys and scoring of immediate mortality and blood-feeding. All live, female *An. gambiae* s.l. were provided access to 10% glucose (w/v) solution and held at ambient conditions in the field laboratory. Delayed mortality was recorded after 24 h for all treatments. Mosquito collections were performed 6 nights per week and on the 7th day, huts were cleaned and aired in preparation for the next rotation cycle.

### Hut trial outcome measures

The efficacy of the experimental hut treatments was expressed in terms of the following outcome measures:

1. Deterrence (%) – proportional reduction in the number of mosquitoes collected in the treated hut relative to the number collected in the control.
2. Exophily (%) – exiting rates due to potential irritant effect of the treatment expressed as the proportion of mosquitoes collected in the veranda trap.
3. Blood-feeding inhibition (%) – proportional reduction in blood-feeding in the treated hut relative to the control. Calculated as follows:

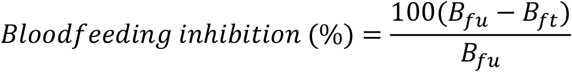

Where *Bfu* is the number of blood-fed mosquitoes in the untreated net control huts and *Bt* is the number of mosquitoes in the huts with insecticide treatments.
4. Mortality (%) – proportion of dead mosquitoes 24 hours after collection.
5. Personal protection (%) – reduction in the number of blood-fed mosquitoes in the treated hut relative to the untreated net control. Calculated as follows:

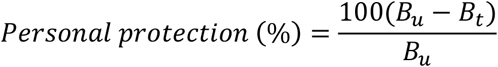

Where *Bu* is the number of blood-fed mosquitoes in the untreated net control huts and *Bt* is the number of mosquitoes in the huts with insecticide treatments.

### Tunnel tests

The tunnel test consists of a square glass cylinder (25 cm high, 25 cm wide, 60 cm in length) divided into two sections using a netting frame fitted into a slot across the tunnel. In one of the sections, a guinea pig was housed unconstrained in a small cage, and in the other section, ~100 unfed female mosquitoes aged 5–8 days were released at dusk and left overnight. The net samples were deliberately holed with nine 1-cm holes to give opportunity for mosquitoes to penetrate the animal baited chamber for a blood meal; an untreated net sample served as the control. The tunnels were kept overnight in a dark room at 25–29oC and 65–85% RH. The next morning, the numbers of mosquitoes found alive or dead, fed or unfed, in each section were scored. Live mosquitoes were provided with 10% glucose solution and delayed mortality was recorded after 24 hours. The pyrethroid-PBO ITNs were compared to Olyset^®^ Net (a permethrin-only net, Sumitomo chemical) and PermaNet^®^ 2.0 (a deltamethrin-only net, Vestergaard Sarl) and an untreated control net. Two to three net pieces were tested per net type.

### Wall cone bioassays

The laboratory-maintained, susceptible *An. gambiae* s.s. Kisumu strain and wild pyrethroid-resistant *An. gambiae* s.l. Covè mosquitoes (F1) derived from breeding sites at the hut station were used for this purpose. At each time point, five cones were attached to the walls and ceiling of the IRS-treated huts. Approximately 50, 3–5-day old mosquitoes were transferred into cones in 5 batches of 10 and exposed to the treated surfaces for 30 mins. As a control, mosquitoes were exposed in cones attached to the walls and ceiling of an unsprayed hut. At the end of exposure, mosquitoes were transferred to netted, plastic cups. Mosquitoes were provided access to 10% (w/v) glucose solution and delayed mortality after 24 h for all treatments.

### Ethical considerations

Ethical approval for the conduct of the study was obtained from the ethics review boards of the Beninese Ministry of Health (Avis éthique No. 34) and the London School of Hygiene & Tropical Medicine (Ethics Ref: 16969). All human volunteer sleepers gave informed written consent prior to their participation; where necessary, the consent form and information sheet were explained in their local language. They were offered a free course of chemoprophylaxis spanning the duration of the trial and up to 3 weeks following its completion. A stand-by nurse was available for the duration of the trial to assess any cases of fever or adverse reactions to test items. Any confirmed cases of malaria were treated free of charge at a local health facility. Animals used as baits in tunnel tests were maintained following institutional standard operating procedures (SOPs) designed to improve care and protect animals used for experimentation. All studies were performed according to relevant national and international guidelines.

### Data analysis

Differences in proportional outcomes (exophily, blood-feeding, mortality) between experimental hut treatments were analysed using a blocked logistic regression model whereas differences in numerical outcomes (mosquito entry) were assessed using a negative binomial regression model. In addition to the fixed effect of the treatment, both models were adjusted to account for variation due to the differential attractiveness of the volunteer sleepers and huts. Results from the different experimental hut trials involving Olyset^®^ Plus and PermaNet^®^ 3.0 were analysed separately. Susceptibility bioassay results were interpreted according to WHO criteria [49]. All analyses were performed in Stata version 15.1.

## Declarations

### Consent for publication

Not applicable

### Authors’ contributions

TS supervised the experiments, analysed the data, and contributed to manuscript preparation. MG, DO, AA and AF conducted the experimental hut trials and tunnel tests. DT performed the resistance bioassays. CN designed the study, supervised the project, and prepared the final manuscript.

## Acknowledgements

We thank the technical staff of CREC (Abibath Odjo, Josias Fagbohoun, Estelle Vigninou, Boris Ndombidje etc) for their assistance. We are grateful to the rice farmers at Covè for their support in the hut study.

## Authors’ information

Not applicable

## List of abbreviations

IRS: Indoor Residual Spraying
WHO: World Health Organization
PQ: Prequalification team
PBO: Piperonyl Butoxide
ITN: Insecticide treated net
LLIN: Long-lasting Insecticidal net
GPIRM: Global Plan for Insecticide Resistance Management
WHOPES: WHO Pesticide Evaluation Scheme
CREC: Centre de Recherche Entomologique de Cotonou
LSHTM: London School of Hygiene & Tropical Medicine
Kdr: Knockdown resistance

## Supplementary information

**Table S1:** Detailed results from tunnel tests with pyrethroid-PBO and pyrethroid-only nets.

| Strain | Susceptible *An. gambiae* Kisumu |  |  |  |  |  | Pyrethroid-resistant *An gambiae* s.l. Covè |  |  |  |  |  |
| --- | --- | --- | --- | --- | --- | --- | --- | --- | --- | --- | --- | --- |
| Net type | Untreated net | Olyset Net | PermaNet 2.0 | Olyset Plus | PermaNet 3.0 (sides) | PermaNet 3.0 (roof) | Untreated net | Olyset Net | PermaNet 2.0 | Olyset Plus | PermaNet 3.0 (sides) | PermaNet 3.0 (roof) |
| N exposed | 307 | 191 | 218 | 191 | 196 | 229 | 208 | 300 | 216 | 303 | 184 | 200 |
| N pass | 236 | 65 | 44 | 11 | 58 | 109 | 107 | 69 | 59 | 53 | 103 | 108 |
| % pass | 76.9 | 34.0 | 20.2 | 5.8 | 29.6 | 47.6 | 51.4 | 23.0 | 27.3 | 17.5 | 56.0 | 54.0 |
| 95% CIs range | 72.2-81.6 | 27.3-40.8 | 14.9-25.5 | 2.5-9.1 | 23.2-36 | 41.1-54.1 | 44.6-58.2 | 18.2-27.8 | 21.4-33.3 | 13.2-21.8 | 48.8-63.2 | 47.1-60.9 |
| N blood fed | 246.0 | 105.0 | 0.0 | 0.0 | 2.0 | 7.0 | 191.0 | 22.0 | 26.0 | 6.0 | 73.0 | 68.0 |
| % blood fed | 80.1 | 55.0 | 0.0 | 0.0 | 1.0 | 3.1 | 91.8 | 7.3 | 12.0 | 2.0 | 39.7 | 34.0 |
| 95% CIs range | 75.7-84.6 | 47.9-62 | 0.0-5 | 0.0-5 | 0-2.4 | 0.8-5.3 | 88.1-95.6 | 4.4-10.3 | 7.7-16.4 | 0.4-3.6 | 32.6-46.7 | 27.4-40.6 |
| % blood-feeding inhibition | - | 31.4 | 100.0 | 100.0 | 98.7 | 96.2 | - | 92.0 | 86.9 | 97.8 | 56.8 | 63.0 |
| N dead 24 h | 34.0 | 187.0 | 218.0 | 190.0 | 196.0 | 227.0 | 5.0 | 125.0 | 58.0 | 272.0 | 62.0 | 136.0 |
| % dead 24 h | 11.1 | 97.9 | 100.0 | 99.5 | 100.0 | 99.1 | 2.4 | 41.7 | 26.9 | 89.8 | 33.7 | 68.0 |
| 95% CIs range | 7.6-14.6 | 95.9-99.9 | 95-100.0 | 98.5-100.0 | 95-100.0 | 97.9-100.0 | 0.3-4.5 | 36.1-47.3 | 20.9-32.8 | 86.4-93.2 | 26.8-40.5 | 61.5-74.5 |

